# A flow cytometry-based assay to quantify the binding of transmembrane ligands to their cognate receptors using fluorescent virus-like particles

**DOI:** 10.64898/2026.05.14.725198

**Authors:** Colin Kim, Maira Gaballa, Danyel Lee, Emmanuelle Jouanguy, Shen-Ying Zhang, Jean-Laurent Casanova, Ahmad Yatim

## Abstract

The binding of transmembrane (TM) ligands to their cognate TM receptors on neighboring cells governs intercellular adhesion and direct cell–cell communication. However, these interactions are difficult to study in vitro because they depend on membrane presentation, ligand orientation, receptor clustering, and avidity, features often not captured by soluble recombinant ligands or cell-free assays. Here, we describe a flow cytometry–based assay using fluorescent, lentiviral-derived virus-like particles (VLPs) displaying TM ligands to quantify binding to their receptors on target cells. Fluorescent VLPs are generated in-house by plasmid transfection in HEK293T cells and enable direct fluorescent detection without fluorochrome-conjugated secondary antibodies. The system is modular and readily accommodates engineered ligand constructs, including patient-derived variants. We applied this platform to generate ICAM-1–displaying fluorescent VLPs and to study human LFA-1 function in patient-derived leukocytes. This protocol provides a detailed workflow for VLP production and in vitro binding assays, offering a simple, quantitative, and cost-effective approach for studying TM ligand–receptor interactions in a membrane context. The system is well suited for mechanistic studies, functional assessment of patient-derived variants, and direct binding assays using patient-derived cells. Integrating the assay into multicolor flow cytometry panels enables simultaneous immunophenotyping and quantification of up to four ligand-receptor interactions at single-cell resolution.

**Key features:**

- Quantifies TM ligand–receptor binding in a membrane context using fluorescent VLPs and flow cytometry.
- Fully in-house, modular system based on plasmid transfection in HEK293T cells, without reliance on recombinant ligands or fluorochrome-conjugated secondary antibodies.
- Supports testing of engineered ligand variants, including patient-derived alleles, and direct functional studies on patient-derived cells.
- Compatible with multicolor flow cytometry panels, enabling simultaneous immunophenotyping and quantification of up to four ligand-receptor interactions at single-cell resolution.

**Graphical overview:** 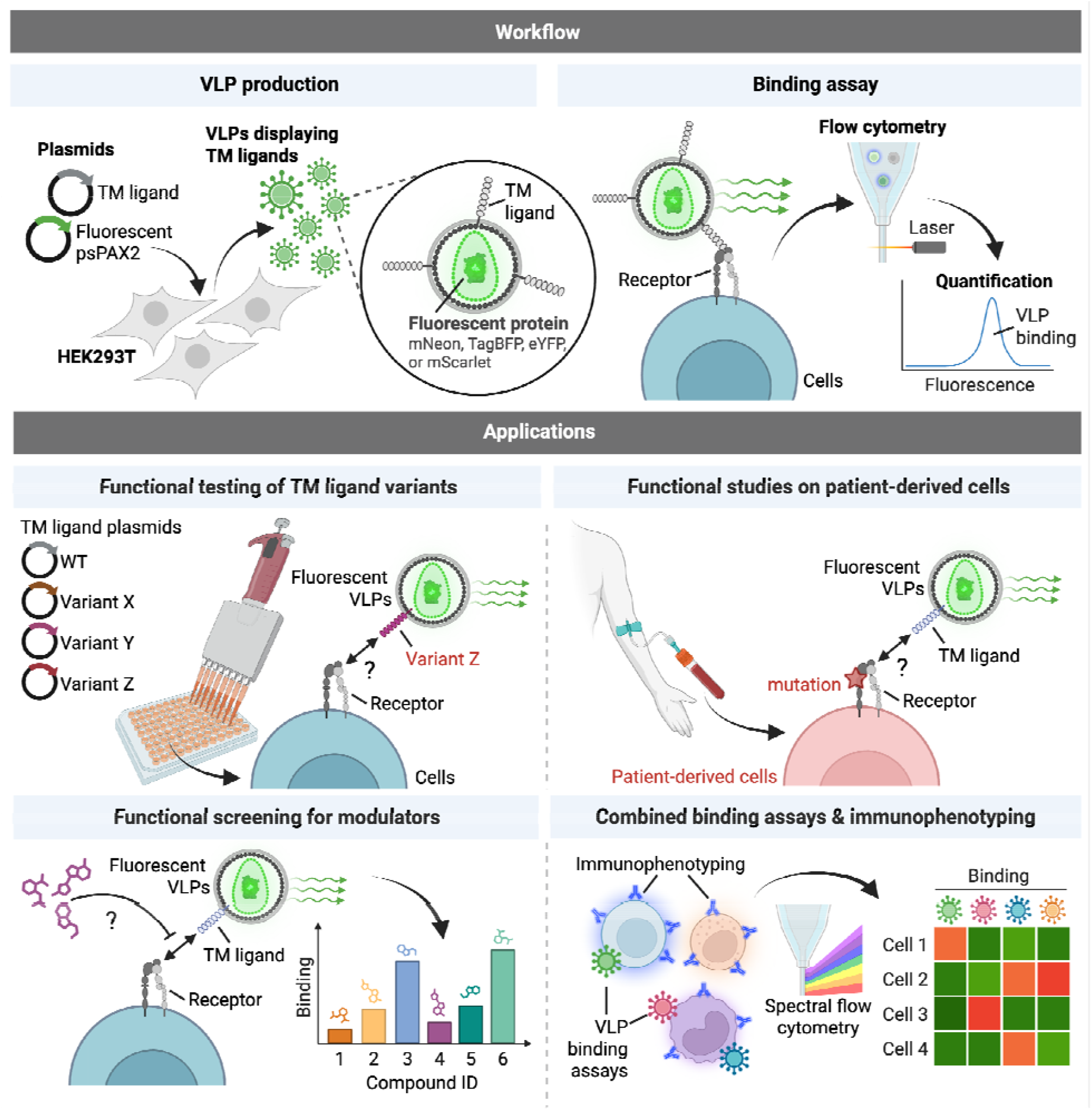

## Background

Transmembrane (TM) ligand–receptor interactions are essential mediators of direct cell–cell communication. Their membrane-bound, contact-dependent nature underlies fundamental biological processes, ranging from tissue patterning and morphogenesis to leukocyte trafficking and activation. For example, leukocyte integrins such as LFA-1 (αLβ2), VLA-4 (α4β1) and α4β7 mediate transendothelial migration through binding to endothelial TM ligands ICAM-1, VCAM-1 and MAdCAM-1, respectively (1). Likewise, T cell receptors (TCRs) and co-stimulatory receptors such as CD28, OX40, 4-1BB or ICOS must engage their cognate TM ligands on antigen-presenting cells for productive T cell priming (2). Accordingly, genetic variants in genes encoding TM ligands or their receptors underlie a broad range of inherited human diseases, including congenital developmental disorders (3, 4) and inborn errors of immunity (5). Notable examples include defects in key lymphocyte costimulatory and coinhibitory molecules—such as CD28 (6), ICOS/ICOSL (7-9), CTLA-4 (10, 11), PD-1/PD-L1 (12-14), OX40 (15), 4-1BB (16), CD40/CD40L (17-21), and CD27/CD70 (22)—as well as essential leukocyte adhesion molecules, such as LFA-1 (1) and other β2-integrins (23-25), highlighting the need for simple assays to evaluate the functional consequences of patient variants in a membrane context. Yet TM ligand-receptor interactions remain challenging to study because they depend on membrane presentation, ligand orientation, receptor clustering, and avidity—features often poorly captured by soluble recombinant proteins or cell-free assays, while cell-based coculture assays can be more difficult to quantify, scale, and standardize. As a result, there is a need for experimental systems that preserve the membrane-associated nature of these interactions while remaining quantitative, scalable, and accessible.

Here, we describe a flow cytometry–based assay using fluorescent virus-like particles (VLPs) displaying TM ligands to quantify receptor binding on target cells. Building on a previously reported lentiviral packaging vector (psPAX2) encoding mNeon fused to the nucleocapsid protein (26), we generated additional psPAX2-derived constructs carrying distinct fluorescent proteins, enabling the production of VLPs labeled with mNeon (Green), eYFP (Yellow), mScarlet (Red), or TagBFP (Blue) and displaying a TM ligand of interest. This platform combines the advantages of membrane-context presentation, experimental flexibility, and robust flow cytometric readout, without the need for fluorochrome-conjugated secondary antibodies. Because VLPs are generated in-house by plasmid transfection in HEK293T cells, the method does not depend on the availability of commercial recombinant ligands and can be implemented in a cost-effective and highly modular manner. Moreover, ligand constructs can be readily engineered to introduce point mutations, domain swaps, or other modifications, making this system particularly well suited for mapping structure–function relationships, assessing the impact of patient variants, and performing direct binding studies on patient-derived cells. We successfully applied this platform to study human LFA-1 using fluorescent VLPs displaying its ligand ICAM-1. The assay enabled quantitative analysis of LFA-1 function on patient-derived leukocytes and contributed to the characterization of a novel inborn error of immunity caused by selective LFA-1 deficiency due to loss-of-function variants in its αL subunit (1).

## Materials and reagents

### Biological materials

1. HEK 293T cells (RRID:CVCL_0063)
2. Jurkat cells (RRID:CVCL_0367)
3. THP-1 cells (RRID:CVCL_0006)

### Plasmids

1. psPAX2-D64V-NC-mNeon (Addgene, Plasmid #196509, a gift from Howard Chang)
2. psPAX2-D64V-NC-mScarlet (This study; to be deposited to Addgene)
3. psPAX2-D64V-NC-TagBFP (This study; to be deposited to Addgene)
4. psPAX2-D64V-NC-eYFP (This study; to be deposited to Addgene)
5. pCMV6-ICAM1 (to be deposited to Addgene)
6. pCMV6-ICAM1 E34A (to be deposited to Addgene)

### Reagents

1. OmniPur BSA, Fraction V (Sigma-Aldrich, catalog number: 2960-500GM), store at 4°C
2. XtremeGENE-9 Transfection Reagent (Roche, catalog number: 6365787001), store at 4°C
3. Lenti-X Concentrator (Takara Biosciences, catalog number: 631231), store at 4°C
4. Fetal Bovine Serum (FBS) (Gibco), store at -20°C Note: FBS should be heat-inactivated at 56°C for 30 min and sterile-filtered through a 0.22 µm filter before use.
5. DMEM + GlutaMAX (Gibco, catalog number: 10566-016), store at 4°C
6. RPMI + GlutaMAX (Gibco, catalog number: 61870-036), store at 4°C
7. OptiMEM Serum-Free Medium (Thermo Fisher Scientific, catalog number: 31985-070), store at 4°C
8. 1M HEPES Buffer (Gibco, catalog number: 15630080), store at 4°C
9. Phosphate Buffered Saline (Corning, catalog number: 21-031-CV), store at room temperature
10. 1M MgCl_2_ (Ambion, catalog number: AM9530G), store at room temperature
11. 0.5M EGTA (bioWORLD, catalog number: 40520008-1), store at room temperature
12. Trypsin EDTA (Gibco, catalog number: 25200-056), store at 4°C
13. Paraformaldehyde (PFA) solution 4% in PBS (ChemCruz, catalog number: sc-281692), store at 4°C and protect from light.

### Solutions

1. DMEM complete medium (see Recipes)
2. RPMI complete medium (see Recipes)
3. Binding buffer (see Recipes)
4. 2X Mg^2+^/EGTA solution (see Recipes)
5. FACS buffer (see Recipes)

### Recipes

#### 1. DMEM complete medium

| Reagent | Final concentration | Quantity or Volume |
| --- | --- | --- |
| DMEM + GlutaMAX |  | 445 mL |
| Heat-inactivated, filtered FBS | 10% v/v | 50 mL |
| HEPES 1M | 10mM | 5 mL |
| Total |  | 500 mL |

#### 2. RPMI complete medium

| Reagent | Final concentration | Quantity or Volume |
| --- | --- | --- |
| RPMI + GlutaMAX |  | 445 mL |
| Heat-inactivated, filtered FBS | 10% v/v | 50 mL |
| HEPES 1M | 10mM | 5 mL |
| Total |  | 500 mL |

#### 3. Binding buffer (RPMI + 0.1% BSA w/v)

| Reagent | Final concentration | Quantity or Volume |
| --- | --- | --- |
| RPMI + GlutaMAX (without FBS) |  | 50 mL |
| BSA Fraction V | 0.1% w/v | 50 mg |
| Total |  | 50 mL |

Combine all components in a 50 mL conical tube and mix by inversion until solid BSA is dissolved. Sterile-filter the solution using a 0.22 µm vacuum filtration unit. Store at 4°C.

#### 4. 2X Mg^2+^/EGTA solution

| Reagent | Final concentration | Quantity or Volume |
| --- | --- | --- |
| Binding buffer |  | 9.82 mL |
| MgCl<sub>2</sub> 1M | 10mM | 100 µL |
| EGTA 0.5M | 4mM | 80 µL |
| Total |  | 10 mL |

Combine all components in a 15 mL conical tube and mix by inversion. Sterile-filter the solution using a 0.22 µm vacuum filtration unit. Store at 4°C.

#### 5. FACS Buffer (PBS + BSA 0.2% w/v)

| Reagent | Final concentration | Quantity or Volume |
| --- | --- | --- |
| PBS |  | 500 mL |
| BSA Fraction V | 0.2% w/v | 1 g |
| Total |  | 500 mL |

Combine all components and mix by inversion until solid BSA is dissolved. Sterile-filter the solution using a 0.22 µm vacuum filtration unit. Store at 4°C.

### Laboratory supplies

1. 10 cm Tissue Culture-Treated Dishes (Falcon, catalog number: 353003)
2. Vacuum filtration unit, Nalgene Rapid-Flow, 0.22 µm, 50 mL (Thermo Fisher Scientific, catalog number: 564-0020)
3. Vacuum filtration unit, Nalgene Rapid-Flow, 0.22 µm, 500 mL (Thermo Fisher Scientific, catalog number: 566-0020)
4. Microcentrifuge tubes, low-binding, 1.5 mL, autoclaved before use (VWR, catalog number: 76332-068)
5. Conical centrifuge tubes, polypropylene, 15 mL, 50 mL (Falcon, catalog number: 352096, 352070)
6. Syringes with luer lock, 10 mL (BD, catalog number: 302995)
7. Acrodisc syringe filters, 0.45 µm (Pall, catalog number: 4614)
8. Serological pipettes, individually wrapped, 5 mL, 10 mL (Falcon, catalog number: 356543, 356551)
9. 96-well polypropylene V-bottom plates (Greiner, catalog number: 651201)

### Equipment

1. Single-channel pipettes 1–10 µL, 20–200 µL, 100–1000 µL
2. Multichannel pipettes 1–50 µL, 20–300 µL
3. Serological pipette controller
4. Hemocytometer or automated cell counter Countess II FL (Thermo Fisher Scientific, catalog number: AMQAF1000)
5. Class II biological safety cabinet
6. Humidified tissue culture incubator, 37 °C and 5% CO_2_
7. Centrifuge with temperature control and speeds up to 1500×g
8. Flow cytometer: Attune NxT (Thermo Fisher Scientific, RRID:SCR_019590) equipped with violet (405 nm), blue (488 nm), and yellow (561 nm) lasers.

### Software and datasets

1. FlowJo Software (v10.10.0, RRID:SCR_008520); requires a license.

## Procedure

### A. ICAM-1-VLP Production

This section describes the production of ICAM-1-displaying VLPs by transfection of HEK293T cells (Figure 1). Perform all cell culture steps using standard aseptic technique. Use high-quality, endotoxin-free plasmid preparations validated by agarose gel electrophoresis and sequencing. To generate VLPs displaying another TM protein of interest, replace the pCMV6-ICAM-1 expression plasmid with the corresponding expression vector (see General note #3).

**Figure 1.**
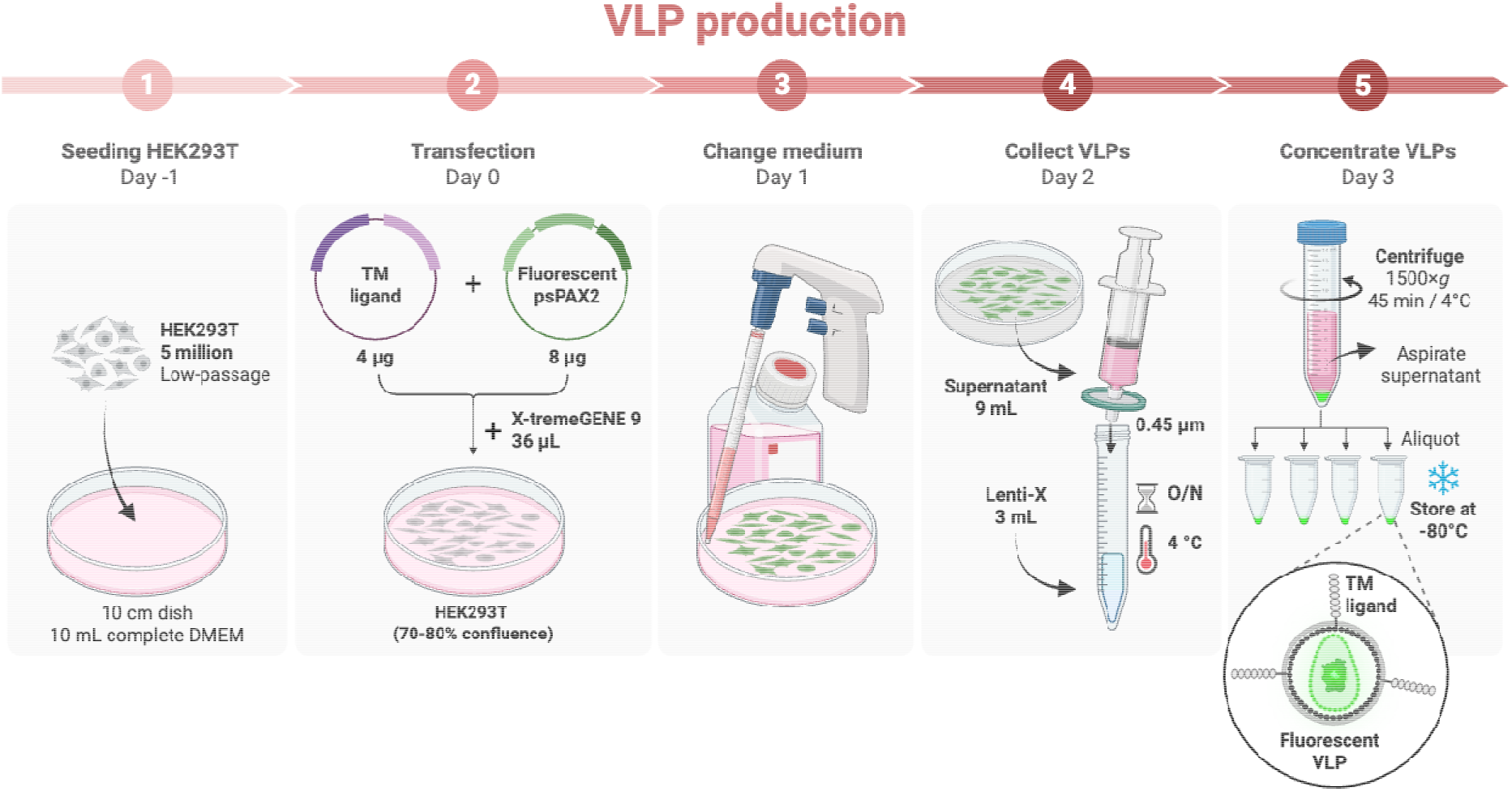
Workflow for production of fluorescent VLPs displaying a TM ligand of interest.

1. Seed 5 million HEK293T cells in 10 mL of complete DMEM per 10 cm tissue culture-treated dish. **Critical:** Ensure homogeneous cell seeding across the dish, as uneven seeding can reduce transfection efficiency. **Note:** All volumes are given for one 10 cm dish. For 6-well plates, scale volumes down fivefold.
2. Incubate the cells overnight at 37 °C in a humidified 5% CO□ incubator.
3. On the following day, confirm that the cells are at 70%–80% confluence and appear healthy, adherent, and evenly distributed, with minimal floating cells or clumping. **Critical:** Do not transfect cells that are overconfluent (>95%), as transfection efficiency may decrease markedly.
4. Prepare the plasmid mix. For each condition, add 8 µg fluorescent psPAX2 plasmid (mNeon, TagBFP, eYFP, or mScarlet) and 4 µg TM ligand expression plasmid (pCMV6-ICAM-1) to 500 µL Opti-MEM in a 1.5 mL low-binding microcentrifuge tube. Mix thoroughly by pipetting.
5. Prepare the XtremeGENE-9 mix. For each condition, add 500 µL Opti-MEM to a 1.5 mL low-binding microcentrifuge tube, then add 36 µL X-tremeGENE 9. Mix thoroughly by pipetting. **Note:** XtremeGENE-9 reagent contains 80% ethanol. Dispense directly into the Opti-MEM without touching the plastic and close the vial immediately after use.
6. Add 500 µL of the X-tremeGENE 9 mix to the plasmid mix for a final transfection volume of 1 mL per condition. Mix thoroughly by pipetting.
7. Incubate for 15 min at room temperature.
8. Add the entire 1 mL transfection mix dropwise to the 10 cm dish of HEK293T cells. **Critical:** Distribute the drops evenly across the dish to ensure uniform distribution.
9. Incubate the transfected cells overnight at 37 °C in a humidified 5% CO□ incubator.
10. After 16-24 h (post-transfection), replace the medium with 10 mL fresh complete DMEM medium per dish. **Critical:** HEK293T cells can detach easily if handled roughly, especially after transfection. Carefully remove the medium from the edge of the dish using a serological pipette or vacuum aspirator, keeping the tip away from the cell monolayer. Then add fresh medium slowly along the wall of the dish to avoid detaching the cells. **Note:** We use complete DMEM containing 10% FBS at this step, which supports robust HEK293T viability and VLP production and did not interfere with downstream ICAM-1–VLP binding assays. Low-serum medium (≤1% FBS) may be used when reducing serum-derived contaminants is important. **Note:** At this stage, transfection efficiency may be assessed by fluorescence microscopy. The fluorescent protein encoded by the psPAX2 construct of choice should be readily detectable in transfected cells. We typically expect >80% positive cells.
11. Incubate the cells for an additional 24 hours at 37 °C in a humidified 5% CO□ incubator.
12. On the following day (48 h post-transfection), harvest the supernatant (∼9 mL) with a sterile 10 mL syringe, then attach a 0.45 µm syringe filter and filter the supernatant into a 15 mL conical tube. **Note:** The supernatant may be collected directly with the syringe by gently tilting the dish and aspirating from the edge, taking care not to touch the cell monolayer.
13. Add 3 mL Lenti-X Concentrator to 9 mL filtered supernatant (1:3, v/v). Mix gently by inversion.
14. Incubate overnight at 4 °C. **Pause point:** According to the manufacturer, samples can be incubated at 4 °C for up to 1 week before centrifugation.
15. On the following day, centrifuge the tubes at 1500×*g* for 45 minutes at 4 °C. **Note:** Pre-cool the tabletop centrifuge to 4 °C before adding the samples.
16. Carefully aspirate the supernatant without disturbing the pellet. **Note:** If residual liquid remains, briefly centrifuge the tubes again at 1,500×g to collect it at the bottom of the tube, then remove it completely with a p200 pipette without disturbing the pellet.
17. Resuspend the pellet in 90 µL of PBS to obtain a 100X VLP stock. **Note:** Mix thoroughly by pipetting until the pellet is fully resuspended, taking care to avoid bubble formation. **Note:** “100X” indicates that the concentrated VLP volume is 100-fold lower than the starting supernatant volume.
18. Use the VLPs immediately or aliquot and store at −80 °C for later use. **Note:** Avoid multiple freeze-thaw cycles.

### B. ICAM-1-VLP Binding Assay

This assay is designed to measure the binding of ICAM-1-displaying VLPs to integrin LFA-1 expressed on target cells. The incubation conditions described here are optimized for broad activation of LFA-1 on THP-1 and Jurkat cells using Mg^2+^/EGTA. Alternative conditions for LFA-1 activation, including other stimuli and cell types, are provided in Table 1. For other TM ligand-receptor pairs, several parameters may require optimization, including the target cell type (which must express the receptor of interest), buffer composition, stimulation conditions, and VLP concentration.

**Table 1.**
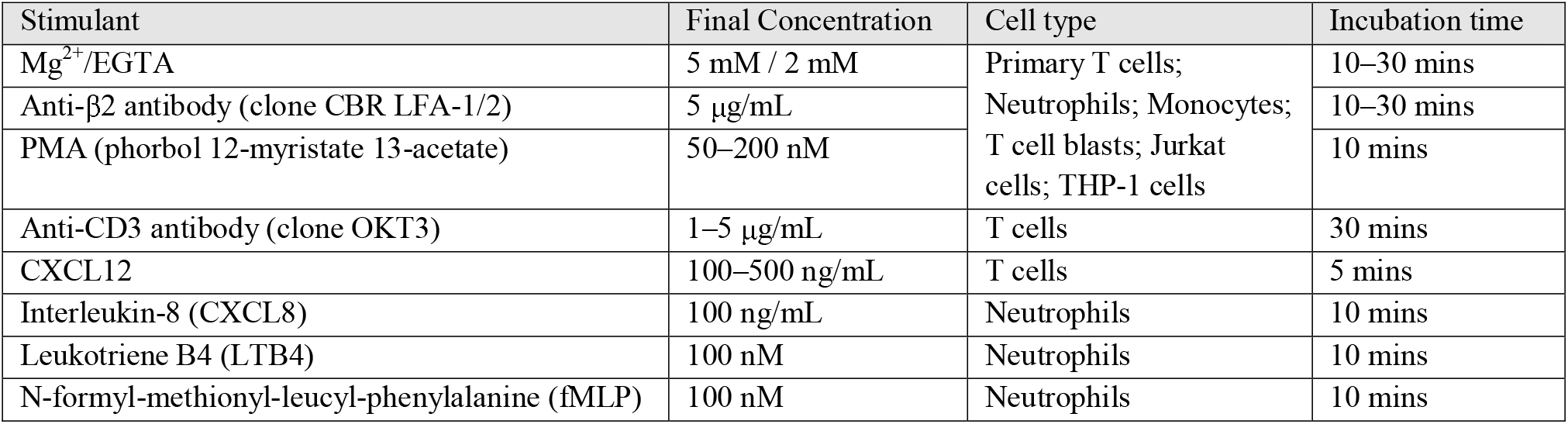
Examples of stimulation conditions for LFA-1 activation in various cell types.

1. Harvest THP-1 or Jurkat cells by centrifugation at 350×g for 5 min, carefully aspirate the supernatant, and resuspend the cell pellet directly in Binding buffer at 1–2 × 10 □ cells/mL. No additional wash step is required before resuspension in Binding buffer. **Note:** THP-1 cells (a monocytic cell line) and Jurkat cells (a T cell line) are non-adherent cells maintained in complete RPMI medium and passaged every 3–4 days at 0.5 × 10□ cells/mL.
2. Distribute 50 µL of cell suspension per well into a 96-well polypropylene V-bottom plate, corresponding to 0.5–1 × 10^5^ cells per well. **Note:** A serum-free binding buffer is recommended for integrin binding assays because it limits background integrin activation, but this may not be necessary for other TM ligand-receptor pairs.
3. Add 50 µL of Mg^2+^/EGTA solution to each well. Mix thoroughly by pipetting. **Note:** This step is used to activate LFA-1. Alternative stimuli for LFA-1 activation may also be used; see Table 1. **Note:** Other TM ligand-receptor pairs may not require an activation step.
4. Immediately add 1–5 μL of concentrated VLP (100X stock solution) to each well. Mix thoroughly by pipetting. **Note:** The optimal VLP amount should be determined by titration of each new preparation and defined empirically as the condition that provides the best specific-to-background signal ratio while remaining below saturating conditions. For ICAM-1-VLPs, we generally obtain excellent results using the 100X stock at a final dilution of 1:100 to 1:20.
5. Incubate at 37 °C for 10 min. **Note:** For ICAM-1-LFA-1 binding, the optimal incubation time depends on the stimulus used (see Table 1) and refers to the time after addition of the activating reagent and VLPs. Incubation time may need to be optimized for other TM ligand-receptor interactions.
6. Add 33 µL of cold 4% PFA to each well to obtain a final concentration of 1%. Mix thoroughly by pipetting and place the plate immediately on ice. Incubate on ice for 20 min. **Note:** Fixation before washing is important for dynamic interactions, such as integrins activated by inside-out signaling (for example, PMA or chemokines; see Table 1), because they can rapidly change conformation and lose ligand binding. For stable TM ligand-receptor interactions, this fixation step may not be required. In these cases, wash steps can be performed at 4 °C before fixation, or fixation may be omitted.
7. Centrifuge plate at 350×*g* for 3 minutes, then remove supernatant. **Note:** The supernatant can be removed by quickly inverting the plate and blotting on a paper towel.
8. Add 250 μL of FACS buffer to each well (Wash #1).
9. Centrifuge plate at 350×*g* for 3 minutes, then remove supernatant.
10. Add 250 μL of FACS buffer to each well (Wash #2).
11. Centrifuge plate at 350×*g* for 3 minutes, then remove supernatant.
12. Add 200 µL FACS buffer to each well to resuspend the cells. Mix thoroughly by pipetting.
13. Acquire the samples on a flow cytometer. **Note:** For Attune NxT, we typically acquire 140 µL of cell suspension per condition using the automated plate sampler. **Pause point:** Fixed and washed cells can be stored at 4 °C for up to 3 days before acquisition.

## Data analysis

### 1. Software and gating strategy

Flow cytometry data can be analyzed using standard analysis software such as FlowJo (BD Biosciences). The following gating hierarchy is recommended to ensure the acquisition of high-quality data: First, exclude debris and gate on the live cell population based on Side Scatter Area (SSC-A) and Forward Scatter Area (FSC-A) profiles, or by the exclusion of a viability dye. Then, exclude doublets or cell aggregates by plotting Forward Scatter Area (FSC-A) against Forward Scatter Height (FSC-H). We recommend acquiring at least 20,000 singlet events per sample; however, lower event numbers are acceptable when analyzing rare cell populations.

### 2. Controls and background subtraction

To ensure accurate quantification of ligand-receptor interactions, the following controls are required: (1) Autofluorescence control: unstained cells used to determine the baseline fluorescence of the target cell population. (2) Non-specific binding control: cells incubated with control VLPs lacking ICAM-1 to assess non-specific or background binding. The gate defining VLP-positive (binding positive) cells should be set using the control VLP lacking ligand (non-specific binding control) and applied consistently across all samples within the same experiment and target cells. As a practical guideline, the positive gate may be positioned so that no more than approximately 1% of cells in the non-specific binding control are scored as positive.

### 3. Quantitative metrics and normalization

Ligand binding should be quantified using the fluorescence intensity carried by the VLPs (mNeon, TagBFP, mScarlet, or eYFP) in the final gated population. Quantification of specific binding can be reported using the Delta Mean Fluorescence Intensity (ΔMFI). This is calculated by subtracting the MFI of the unstained or non-specific binding control from the MFI of the VLPs displaying ICAM-1. To evaluate the effect of treatment or cellular activation on the binding, report the relative binding as the ratio of the ΔMFI of the treated sample to the ΔMFI of the untreated control. To assess the impact of genetic mutations, report the ΔMFI of the mutant ICAM-1 VLP as a percentage or ratio relative to the wild-type (WT) ICAM-1 VLP binding value.

### 4. Statistical Analysis and Reproducibility

To confirm statistical significance, it is recommended that experiments be performed in at least three independent biological replicates. Comparisons between groups (e.g., WT vs. Mutant) should be performed using appropriate statistical tests, such as a two-tailed Student’s t-test or ANOVA with multiple comparisons, as described in (1). No specific computational expertise in Linux or R is required for these standard analyses.

## Validation of protocol

### 1. Validation of fluorescent VLP detection and performance in binding assays

To assess the performance of the assay across different fluorescent proteins, we tested four fluorescent VLP variants (Figure 2A, 2B) spanning a broad range of the visible spectrum: mNeon (excitation: 506 nm, emission: 517 nm), eYFP (excitation: 514 nm, emission: 527 nm), mScarlet (excitation: 569 nm, emission: 594 nm), and TagBFP (excitation: 400 nm, emission: 456 nm) (Figure 2C). These VLPs either displayed ICAM-1 or served as ligand-negative controls generated with the corresponding empty vector (EV). Binding assays were performed using THP-1 cells, which express ICAM-1 receptor LFA-1. Robust and specific binding was observed for all four ICAM-1–displaying VLPs, whereas EV-VLPs showed little to no binding (Figure 2D). Although all fluorescent VLPs enabled reliable detection of binding, the highest fluorescence intensity was obtained with VLPs carrying mNeon, whereas eYFP yielded the lowest relative signal (Figure 2D).

**Figure 2.**
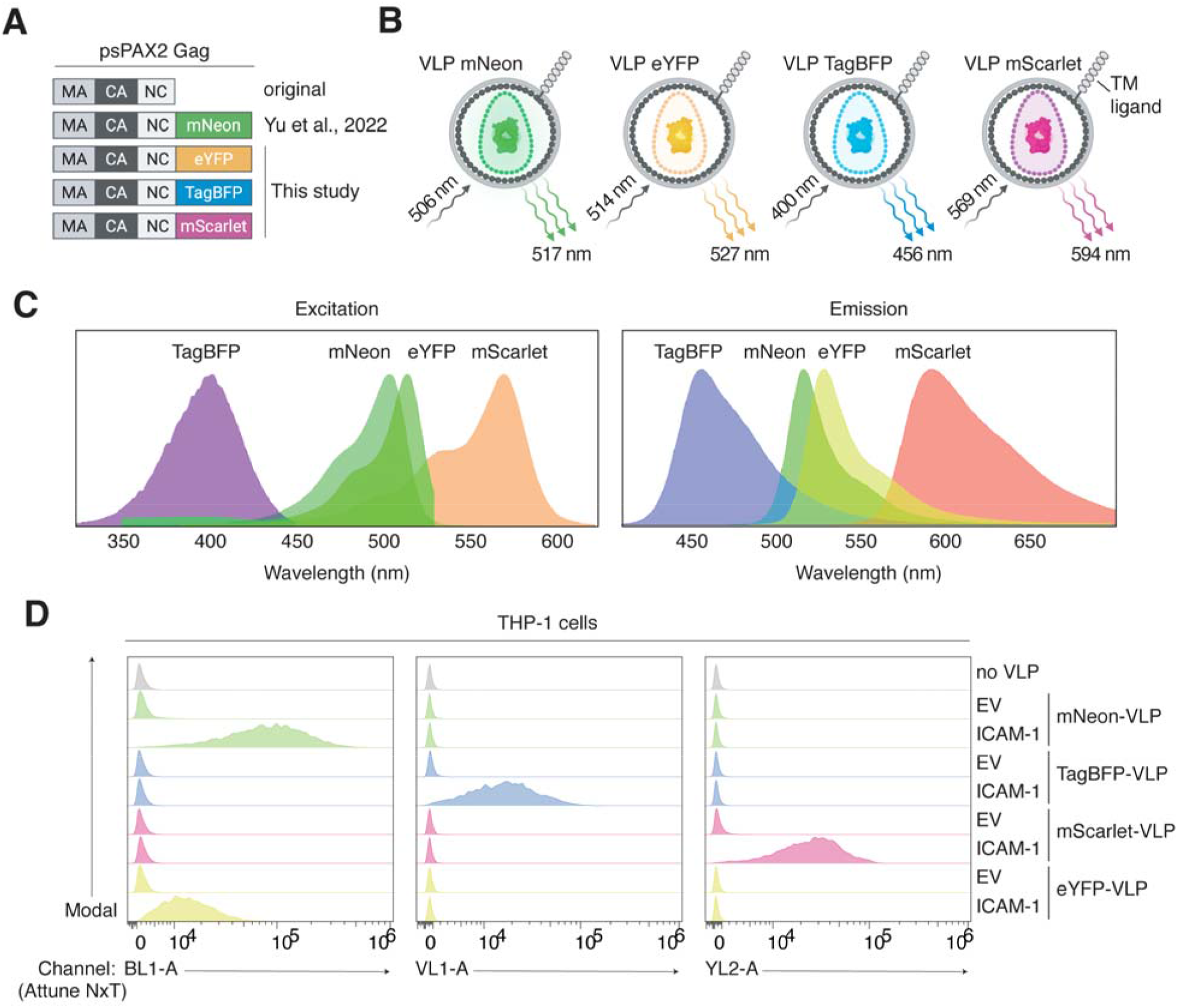
Generation and validation of fluorescent VLPs for TM ligand–receptor binding assays. A. Schematic of the psPAX2 Gag constructs used to generate fluorescent VLPs. B. Schematic of fluorescent VLPs carrying mNeon, eYFP, TagBFP, or mScarlet and displaying a TM ligand at their surface. Approximate excitation and emission maxima are indicated for each fluorescent protein. C. Excitation and emission spectra of mNeon, eYFP, TagBFP, and mScarlet. D. Flow cytometry histograms showing binding of fluorescent VLPs to THP-1 cells. Cells were incubated with VLPs displaying ICAM-1 or generated with empty vector (EV). Histograms are shown for the corresponding fluorescence channel of each VLP type.

### 2. Validation of binding specificity through receptor knockout

The specificity of ICAM-1 VLP binding was validated using CRISPR/Cas9-mediated knockout (KO) of the αL subunit of LFA-1 in Jurkat cells (Figure 3A). We compared the binding of ICAM-1 VLPs to parental Jurkat cells and αL KO Jurkat cells. Deletion of αL resulted in complete loss of surface LFA-1 expression (Figure 3B) and a corresponding complete loss of ICAM-1 VLP binding (Figure 3C). Binding in αL KO cells was reduced to the background level observed with control VLPs lacking ICAM-1 (EV-VLPs) (Figure 3C). These results confirm that the binding (fluorescent signal) is strictly dependent on the expression of the cognate receptor on the target cells. Additional validation data, including quantification, statistical analyses, and the generation of the αL KO Jurkat cells, are provided in Yatim et al., 2026 (1).

**Figure 3.**
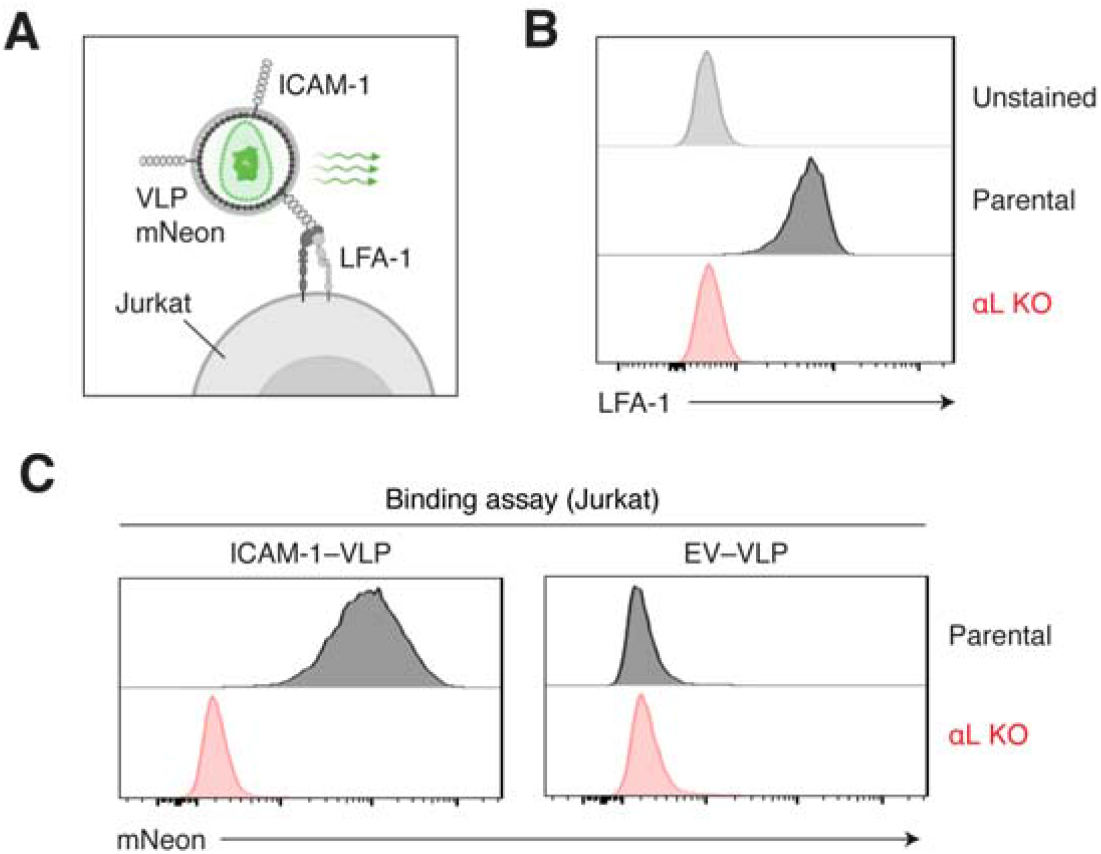
Validation of binding specificity using αL knockout Jurkat cells. A. Schematic of the binding assay using VLPs displaying ICAM-1 to detect binding to LFA-1 on Jurkat cells. B. Flow cytometry histograms showing surface LFA-1 expression in parental and αL knockout (αL KO) Jurkat cells, assessed with the TS2/4 clone. Unstained Jurkat cells are shown as the background autofluorescence control. C. Representative flow cytometry histograms showing binding of ICAM-1– displaying VLPs (ICAM-1–VLPs) or control VLPs lacking ICAM-1 (EV-VLPs) to parental and αL KO Jurkat cells.

### 3. Functional testing of TM ligand variants

To test the ability of the assay to detect loss-of-function ligand variants, we introduced the E34A substitution into the ICAM-1 construct by site-directed mutagenesis, as previously described (1). This residue is critical for the interaction between ICAM-1 and the αL subunit of LFA-1 (27, 28). VLPs displaying wild-type ICAM-1 or the ICAM-1-E34A variant were produced and tested in binding assays on Jurkat cells (Figure 4A). The E34A substitution abolished VLP binding, confirming that this variant is defective for LFA-1 binding (Figure 4B). These results support the use of the assay for structure–function studies and for functional evaluation of patient-derived TM ligand variants. Additional data, quantification, and statistical analyses are provided in Yatim et al., 2026 (1).

**Figure 4.**
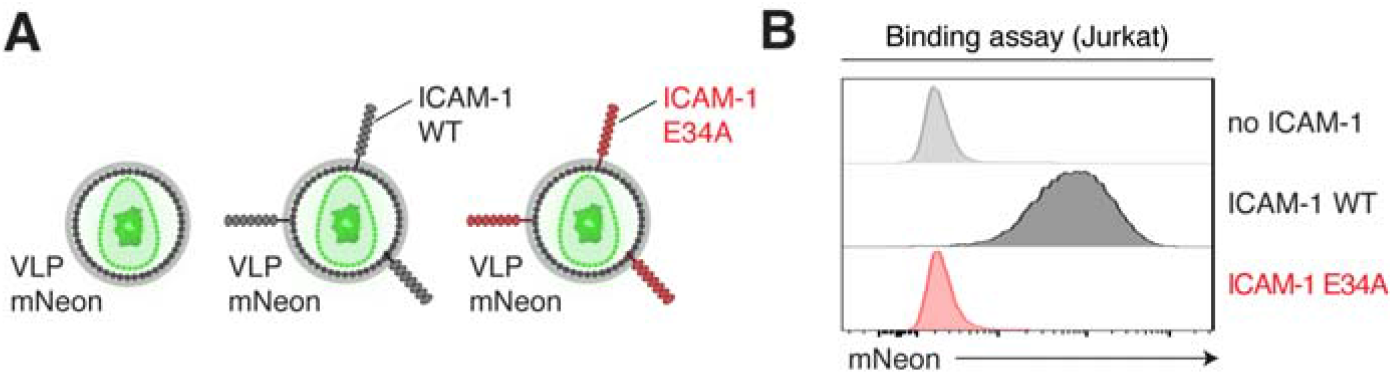
Functional testing of an ICAM-1 variant in the VLP binding assay. A. Schematic of VLPs displaying wild-type (WT) ICAM-1 or the LFA-1-binding–defective ICAM-1 variant Glu34Ala (E34A). B. Representative flow cytometry histograms showing binding of VLPs displaying ICAM-1 WT or ICAM-1-E34A to Jurkat cells.

### 4. Functional assessment of receptor activity in patient-derived cells

We tested binding of ICAM-1 VLPs on T cell blasts generated from healthy donors and from patients harboring biallelic loss-of-function variants in *ITGAL*, the αL subunit of LFA-1. T cell blasts from αL-deficient patients failed to capture VLPs displaying ICAM-1, indicating a complete loss of LFA-1–mediated ICAM-1 binding, consistent with their impaired adhesion to and migration on ICAM-1. These findings show that the VLP binding assay is suitable for functional interrogation of receptor activity in patient-derived cells. Flow cytometry plots, quantification, statistical analyses, and the detailed method for T cell blast generation are available in Yatim et al., 2026 (1), specifically in Figure 3 and the Methods section.

### 5. Integration of the binding assay into high-parameter spectral flow cytometry

To enable simultaneous quantification of ligand binding and deep immunophenotyping at single-cell resolution, the ICAM-1 VLP binding assay was performed using peripheral blood mononuclear cells (PBMCs) and followed by staining with a multicolor spectral flow cytometry panel. VLPs were added concomitantly with the integrin-activation stimulus and incubated as described in the Procedure section. Cells were then fixed to preserve VLP binding, washed, and stained with the antibody panel. Most flow cytometry antibody clones are compatible with fixation before staining, but this should be validated for each antibody. This workflow allowed ligand binding to be assessed across a broad range of leukocyte subsets in response to various integrin-activating stimuli (1). Detailed methods and associated data, including antibody catalog numbers, are provided in Yatim et al., 2026 (1).

## General notes and troubleshooting

### General notes

#### 1. Ligand incorporation into the VLP membrane

This protocol is based on the principle that any TM ligand expressed at the plasma membrane of HEK293T cells after transfection will be incorporated and displayed on the surface of the budding VLPs. While validated here for ICAM-1 and LFA-1, the system is modular and can be adapted to study a wide range of TM ligand–receptor interactions. However, protein-specific motifs can occasionally influence the efficiency of viral incorporation. Since ICAM-1 is known to be enriched in HIV-derived virions (29), other TM proteins may demonstrate lower incorporation propensities. In such cases, incorporation of the TM ligand into purified or concentrated VLPs can be assessed directly by immunoblotting, using an antibody against the ligand or an epitope tag when available. If incorporation is poor, we recommend generating a chimeric construct by fusing the extracellular domain of the protein of interest to the transmembrane and cytoplasmic domains of ICAM-1 or another well-incorporated protein, a strategy successfully employed in previous studies (26) (see General note 3).

#### 2. VLP quantification and normalization

Some applications, such as variant screening or comparison of multiple ligand constructs, may require concentrated VLP preparations to be normalized so that differences in binding reflect ligand properties rather than variation in VLP production. We have successfully quantified HIV-derived VLPs using a p24 ELISA kit (Takara, Cat. No. 631476). In the setting of variant screening, reduced binding may result from reduced expression or stability of the mutant ligand, or from impaired binding activity. Because both are relevant mechanisms of loss-of-function, we do not recommend correcting for mutant expression in the primary screen. In this context, normalization of total VLP input is generally sufficient. When the goal is to distinguish more precisely between loss-of-expression versus loss-of-binding activity, additional normalization is needed. In that case, WT and mutant constructs should be matched for both surface expression on producer HEK293T cells and incorporation into VLPs. This can be achieved by adjusting the amount of TM ligand plasmid used during transfection to obtain similar surface expression by flow cytometry and similar incorporation into purified VLP preparations, for example by immunoblotting.

#### 3. Cloning of TM ligands and design of chimeric constructs

The plasmid encoding human ICAM-1 was constructed by amplifying the ICAM-1 cDNA from Addgene plasmid no. 8632 (RRID: Addgene_8632; a gift from T. Springer) and inserting it into a pCMV6 backbone by In-Fusion cloning (Takara Bio, Cat. No. 638946). More generally, the cDNA encoding the TM ligand of interest, PCR-amplified from cDNA or another expression vector, can be cloned into pCMV6 or another mammalian expression vector using standard restriction enzyme–based cloning or In-Fusion assembly, depending on the available restriction sites and the sequence of the insert. Addition of an epitope tag should be avoided when possible, as it may interfere with ligand folding, receptor binding, or incorporation into VLPs. If a tag is required to facilitate detection by flow cytometry or immunoblotting, its position should be carefully selected and validated to confirm that it does not affect surface expression, VLP incorporation, or binding activity. For TM proteins that are poorly incorporated into VLPs, ICAM-1-based chimeric constructs may be considered (see General note #1). Yu et al. (26) successfully used chimeric constructs to display proteins of interest on lentiviral particles by fusing their extracellular domains to the ICAM-1 transmembrane domain (IVIITVVAAAVIMGTAGLSTYLY) and cytoplasmic tail (NRQRKIKKYRL). The fusion design should preserve the complete extracellular domain of the ligand of interest and then validated for surface expression, VLP incorporation, and receptor binding.

### Troubleshooting

#### Problem 1: Low VLP production or poor HEK293T transfection efficiency

##### Possible cause

Poor plasmid quality; overconfluent HEK293T cells at the time of transfection; high passage number.

##### Solution

Use high-quality, endotoxin-free plasmid preparations (e.g., Midiprep or Maxiprep) and verify integrity via agarose gel electrophoresis and/or sequencing. Ensure HEK293T cells are maintained in exponential growth phase, are mycoplasma-free, and are transfected at ∼70% confluency. Avoid using cells at high passage for VLP production, as transfection efficiency and budding capacity can decline.

#### Problem 2: High background or non-specific binding of control VLPs

##### Possible cause

Excessive VLP concentration; active VLP internalization (e.g., phagocytosis by myeloid cells); receptor-independent binding of VLPs to target cells.

##### Solution

Perform a VLP titration to determine the optimal concentration that maximizes the specific-to-non-specific signal ratio. Include appropriate negative controls, such as receptor-specific knockout cells and empty VLPs produced in the absence of the TM ligand. Shorten the incubation time to limit alternative uptake mechanisms.

#### Problem 3: Weak or absent signal in the binding assay

##### Possible cause

Inefficient VLP production; low surface expression or poor VLP incorporation of the transfected TM ligand; low receptor density on target cells; transient or low-affinity ligand–receptor interaction.

##### Solution

First, confirm HEK293T transfection efficiency by fluorescence microscopy at 24–48 hours post-transfection. Then assess TM ligand surface expression on HEK293T cells by flow cytometry and, when needed, assess ligand incorporation directly in purified or concentrated VLP preparations by immunoblotting. If incorporation is poor, optimize TM ligand incorporation into the VLP membrane (see General note #1). Finally, verify receptor expression on target cells by flow cytometry. Because each VLP contains multiple fluorescent molecules, receptor-dependent binding is expected to be readily detectable for high-affinity/avidity ligand–receptor interactions. For transient or low-affinity interactions, VLP input, incubation time, and binding conditions may need additional optimization.

## Acknowledgments

We thank all members of both branches of the Laboratory of Human Genetics of Infectious Diseases for discussions and technical support; Yelena Nemirovskaya, Mark Woollett and Kerel Francis for administrative assistance; members of the Flow Cytometry Resource Center (RRID:SCR_017694) at The Rockefeller University for technical assistance. The graphical abstract and figure schematics were created with BioRender.

## Author contributions

Conceptualization, C.K. and A.Y.

Investigation, C.K., M.G., D.L. and A.Y.

Writing—Original Draft, C.K. and A.Y.

Writing—Review & Editing, C.K., M.G., D.L., E.J., S.-Y.Z., J.-L.C. and A.Y.

Funding acquisition, E.J., S.-Y.Z., J.-L.C. and A.Y.

Supervision, E.J., S.-Y.Z., and A.Y.

## Original research papers

The protocol was described and validated in (1), using the original fluorescent lentiviral construct developed by (26).

